# The effect of offspring number on the adaptation speed of a polygenic trait

**DOI:** 10.1101/212613

**Authors:** Markus Pfenninger

**Affiliations:** Senckenberg Biodiversity and Climate Research Centre, Frankfurt am Main, Hessen, Germany Molecular Ecology Group, Institute for Ecology, Evolution & Diversity, Goethe-University Frankfurt am Main, Hessen, Germany

## Abstract

There is increasing evidence that rapid phenotypic adaptation of quantitative traits is not uncommon in nature. However, the circumstances under which rapid adaptation of polygenic traits occurs are not yet understood. Building on previous concepts of soft selection, i.e. frequency and density dependent selection, I developed and tested the hypothesis that adaptation speed of a polygenic trait depends on the number of offspring per breeding pair in a randomly mating diploid population.

Using individual based modelling on a range of offspring per parent (2–200) in populations of various size (100–10000 individuals), I could show that the by far largest proportion of variance (42%) was explained by the offspring number, regardless of genetic trait architecture (10–50 loci, different locus contribution distributions). In addition, it was possible to identify the majority of the responsible loci and account for even more of the observed phenotypic change with a moderate population size.

The simulation results suggest that offspring numbers may a crucial factor for the adaptation speed of quantitative loci. Moreover, as large offspring numbers translates to a large phenotypic variance in the offspring of each parental pair, this genetic bet hedging strategy increases the chance to contribute to the next generation in unpredictable environments.

*“Hence, as more individuals are produced than can possibly survive, there must in every case be a struggle for existence, either one individual with another of the same species, or with the individuals of distinct species, or with the physical conditions of life.”* (Darwin, 1859, chapter III)

## Introduction

There is increasing evidence for rapid adaptation of phenotypic traits in nature (see references in [1]). The speed with which adaptation proceeds depends, amongst other issues, on the intensity of selection [2]. In the standard model of population genetics, however, selection coefficients (s) are assumed to be a fixed character of a particular allelic substitution, respectively the genotype. This is increasingly viewed as insufficient to explain observed rates of phenotypic evolution [1]. However, there is increasing evidence that selection coefficients may vary in time and space [3, 4]. In addition, most adaptive events possibly occur by polygenic adaptation, which is qualitatively different from models of adaptive substitutions [5]. Polygenic adaptation allows rapid adaptation by small shifts in allele frequencies and without the need for adaptive fixation events of any locus involved [1]. Such an infinitesimal model of adaptation, acting on a very large number of loci, is widely applied in quantitative genetics to predict phenotypic change in both natural and artificial selection [6].

However, the allele frequency changes at each individual locus are rather small and thus may be indistinguishable from drift [1], which would make the identification of the genomic basis of adaptation difficult. New efforts are therefore necessary to bridge the gap between population genetic theory and empirical data, in particular to develop predictions and hypothesis on rapid polygenic adaptation that can be empirically tested [6].

I want to recur here to ideas that the strength (and thus the speed) of selection on polygenic traits may vary depending on the density of a population. This goes ultimately back to Darwin [7], who identified overproduction as one necessary prerequisite for any natural selection to happen. Expressed in more technical terms, according to Darwin natural selection can thus only happen, if N, the number of individuals produced in a population, is larger than K, the carrying capacity. This implies indirectly that, as the ratio between N and K gets larger, selection should be more intense. In a mathematical approach, MacArthur [8] tried to link differential birth rate of genotypes (i.e. fitness) with the relation between the population size and the carrying capacity K. Roughgarden found indeed a few years later that the strength of natural selection varies with population density such that selection is stronger when density is higher [9]. Wallace [10], by introducing the concept of soft selection, defined it as both density and frequency dependent and could show that this sort of selection is applicable to many realistic ecological scenarios. Based on these theoretical considerations and empirical evidence, Reznick even championed soft selection as the prevailing mode in nature [11]. Charlesworth [12] recently considered soft selection as potential solution for the conflict between certain expectations from theoretical population genetics and empirical data.

Expanding on these ideas, I want to test the hypothesis that differential numbers of offspring per breeding pair influence the speed of adaptation in response to a sudden shift in the optimal fitness mean, everything else being equal. The rationale behind this hypothesis is simple: In a constant sized population, the mean number of offspring per breeding pair determines the degree of overproduction and thus the degree of competition among the offspring. A highly competitive situation arises from a large number of offspring and thus, even small fitness differences may determine whether an individual gets to reproduction or not. Viewed from the perspective of the parents, the more offspring a breeding pair has, the wider the phenotypic range of the offspring at polygenic, i.e. quantitative traits will be and the more evenly the possible trait space as defined by the genotypic composition of the parents will be filled; i.e. the chances of producing offspring with phenotypes close to a new selective optimum will increase with the number of offspring per breeding pair. In other words, the phenotypic range to select the next generation of breeding pairs from is wider and therefore contains more individuals closer to the phenotypic optimum that can be selected. Hence, the trait mean of the next generation is closer to the new optimum – adaptation should proceed faster.

I used an individual based modelling (IBM) approach to evaluate the effect of different offspring numbers on the speed of adaptation of a polygenic trait in response to a sudden shift of the fitness optimum. In addition, the model was used to evaluate the possibility to detect the loci underlying the quantitative trait by comparison of their selection driven allele frequency change with those from neutral markers from before-after data.

## Material & Methods

A population in the simulation consisted of N_b_ breeding adult individuals that randomly mated with another individual to produce 2*N_b_*x_juv_ offspring where x_juv_ stands for the number of offspring per individual. A generation cycle started therefore with 2*N_b_*x_juv_ = N_tot_ individuals of which only N_b_ individuals got to reproduction to create the next generation. The constant sized population thus had non-overlapping, discrete generations. Mutations were not taken into account to keep the number of parameters low and because they likely play a minor role in fast adaptation.

The genomes of individuals consisted of unlinked biallelic loci. The phenotypic trait values in arbitrary units were determined by n biallelic quantitative trait loci (QTL) with strictly additive effects. To evaluate the effect of different genetic architectures, the phenotypic trait contribution of each locus was drawn from a gamma distribution with alpha parameters 0.1 and 1.5. The smaller alpha produces a distribution where all loci contribute almost equally to the trait. With the larger value, few loci contribute most to the phenotypic value (Figure 1). The actual phenotypic contribution values for the two alleles of each locus were then determined by drawing random values from a Gaussian distribution with mean and standard deviation as given by the gamma distribution value for the respective locus. The quantitative trait was influenced by 10, 30 and 50 QTL. This relatively low quantitative trait locus number was used, because i) not so many recombination events may be expected during the few generations expected to achieve adaptation, thus reducing the effective number of truly independent sites in the genome and ii) even if many more loci may contribute to a quantitative trait, only a fraction of them is probably free to vary, because of negative pleiotropic interactions with other loci [5]. Allele frequencies for each quantitative trait locus were drawn from a uniform distribution bounded by 0.1 and 0.9. In addition, 100 neutral biallelic loci (NL) were simulated, with initial allele frequencies drawn from a uniform distribution bounded as before.

**Figure 1.**
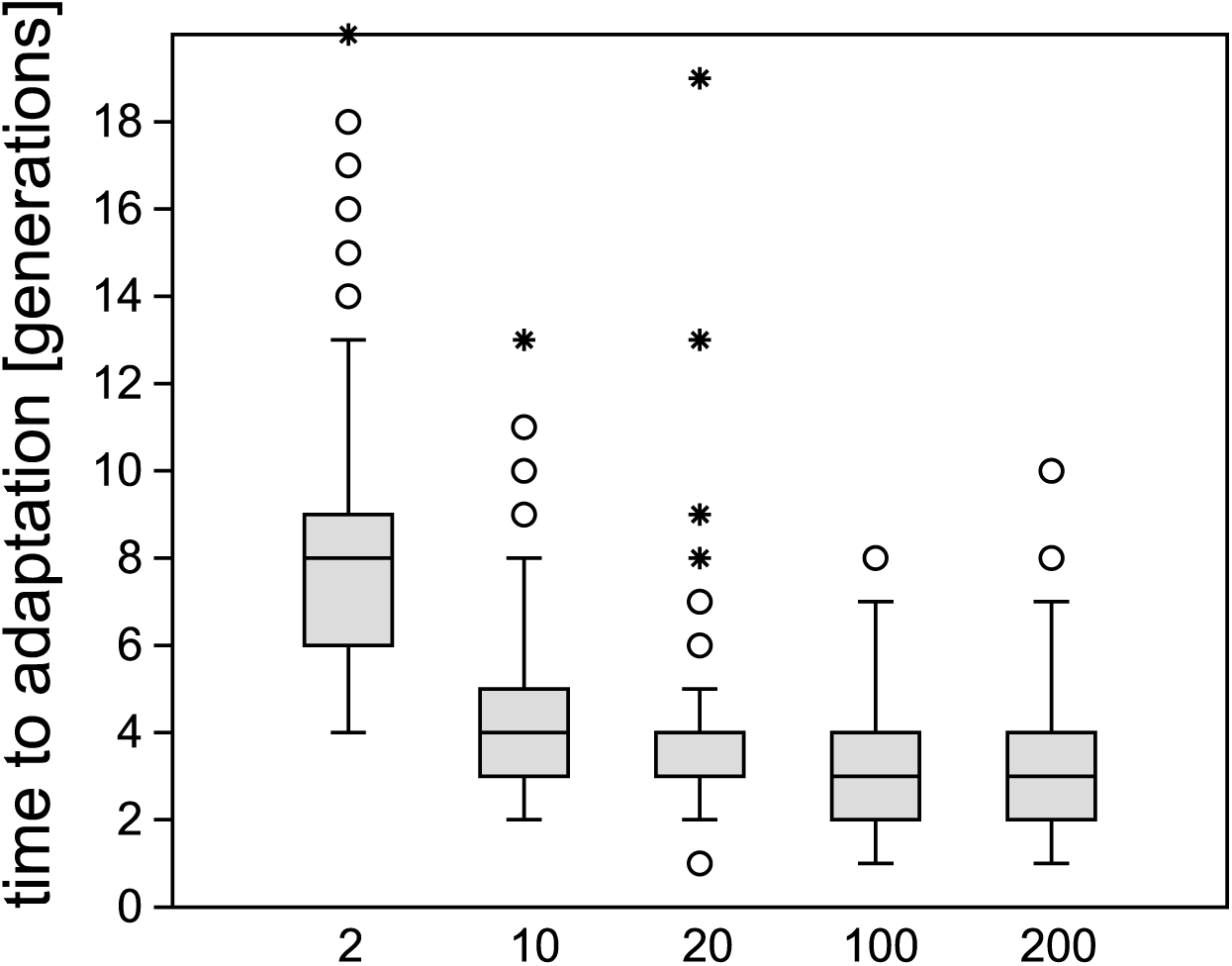
Boxplot of the effect of different numbers of offspring per breeding pair on the time to adaptation.

The initial population started in the juvenile stage, which means that N_b_*x_juv_ were created by randomly assigning two alleles for each locus (QTL and NL) with probabilities according to the respective allele frequency for the locus to each individual. The values of the QTL were then used to calculate the phenotypic value for each individual. Once the entire population was created, the population mean of the trait was determined.

In the first generation, a Gaussian stabilizing selection regime was applied. Individuals were drawn at random with a probability to get to reproduction determined by the deviation of the individual from the population mean until the adult population size was reached. From this adult base population, allele frequencies for QTL and NL were determined, the phenotypic population mean and standard deviation calculated and all values recorded. The new fitness optimum was set to the mean plus three times the standard deviation of the adult base population.

Reproduction took place by randomly drawing two individuals without replacement from the adults of the previous generation. To produce an offspring individual, one random allele from each locus was drawn from each of the parents. This was repeated for each adult pair until 2*N_b_*x_juv_ offspring were created. All offspring from all pairs was first randomly mixed and then sorted according to the individual deviation from the new fitness optimum. All but the N_b_ individuals with the smallest deviation from the new fitness optimum were then deleted. This corresponds to a deterministic selection regime. Allele frequencies for all loci, phenotypic population mean and standard deviation were recorded. This procedure was repeated until the population mean was equal to or larger than the new fitness mean or until adaptation failed because of a lack of genetic variation. The number of generations taken to achieve this shift of the population mean was taken as the speed of adaptation.

For identification of loci responsible for the observed adaptation, I compared the allele frequency shift between the initial adult base population and the generation when adaptation was achieved. The number of loci that went to fixation were recorded for both QTL and NL. A QTL allele frequency shift deemed to be significantly identified if it was larger than those of 99% of NL. It was also recorded for what proportion of the observed phenotypic change the such identified loci were responsible and whether there was a correlation between the observed allele frequency shift and the magnitude of the phenotypic variation caused by the respective locus. The IBM simulation was written in Python 3.4 and run on normal desktop PCs, the source code can be found in the Supplement.

Simulations were run with the parameters given in Table 1 in all possible combinations. Each parameter combination was run with 10 repetitions. The data was analysed after appropriate transformation (proportion variables were arcsin, continuous variables and factors log transformed) with a General Linear Mixed Model in R [13]. N_b_, xj_uv_ and QTL were considered as continuous factors, alpha as categorical factor. Whenever appropriate, the model was simplified.

**Table 1.** Parameters and their values as used in simulations. All parameters were run in all possible combinations with ten replicates each, amounting to 1200 simulations in total.

| Parameter | Explanation | Values used in simulation |
| --- | --- | --- |
| $N_b$ | Number of breeding adults | 10, 100, 1000, 10000 |
| $x_{juv}$ | Number of offspring produced by each parent | 2, 10, 20, 100, 200 |
| QTL | Number of quantitative trait loci determining the phenotypic value of the trait | 10, 30, 50 |
| alpha | Parameter alpha of a gamma distribution determining the contribution distribution of quantitative trait loci | 0.1, 1.5 |

## Results

Over all simulations, it took on average 4.60 generations to shift the phenotypic trait mean three initial standard deviations, with a range of 1 to 38 generations. The differences between adaptation time means among treatments were best explained by the number of offspring per parent, accounting for 42.6% of the variance (Table 2). The mean differences in time to adaptation varied between 8.36 (3.37 s.d.) generations for 2 offspring per parent and 2.98 (1.34 s.d.) and thus almost threefold. The relation seems to converge in a non-linear fashion to a minimum value with increasing offspring numbers (Figure 1) There were also significant trait mean differences for the factors N_e_, number of QTL and alpha treatments, however, taken together, these factors accounted for less than 5% variance (Table 2).

**Table 2.**
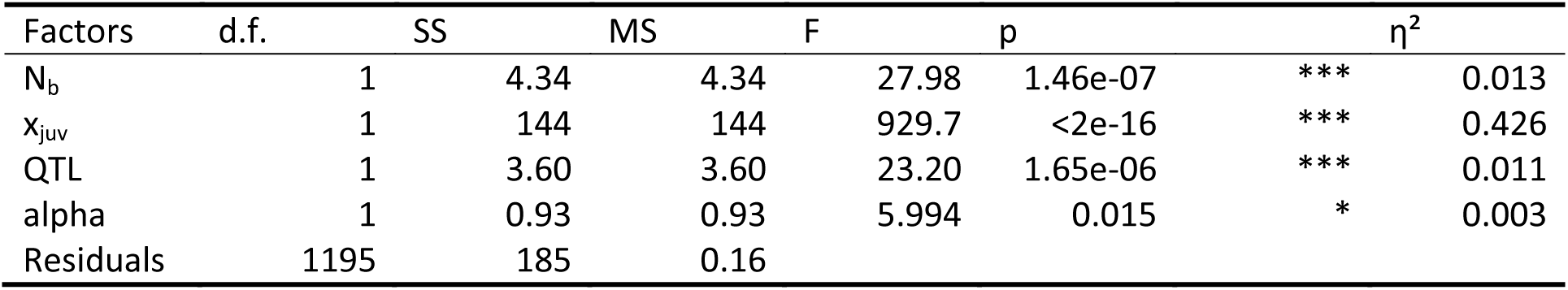
Simplified GLM to explain variance in adaptation time.

On average, about one fifth of all QTL went to fixation (0.213). In individual simulations, the entire range from none to all QTL fixed was observed. A mean of 0.061 NL went to fixation, at most, 60% of NL loci fixed in any single run. There was a strong correlation (0.757) between the phenotypic effect of a locus and the observed allele frequency shift, however, in individual runs, also no or even negative correlations were observed (Figure 2).

**Figure 2.**
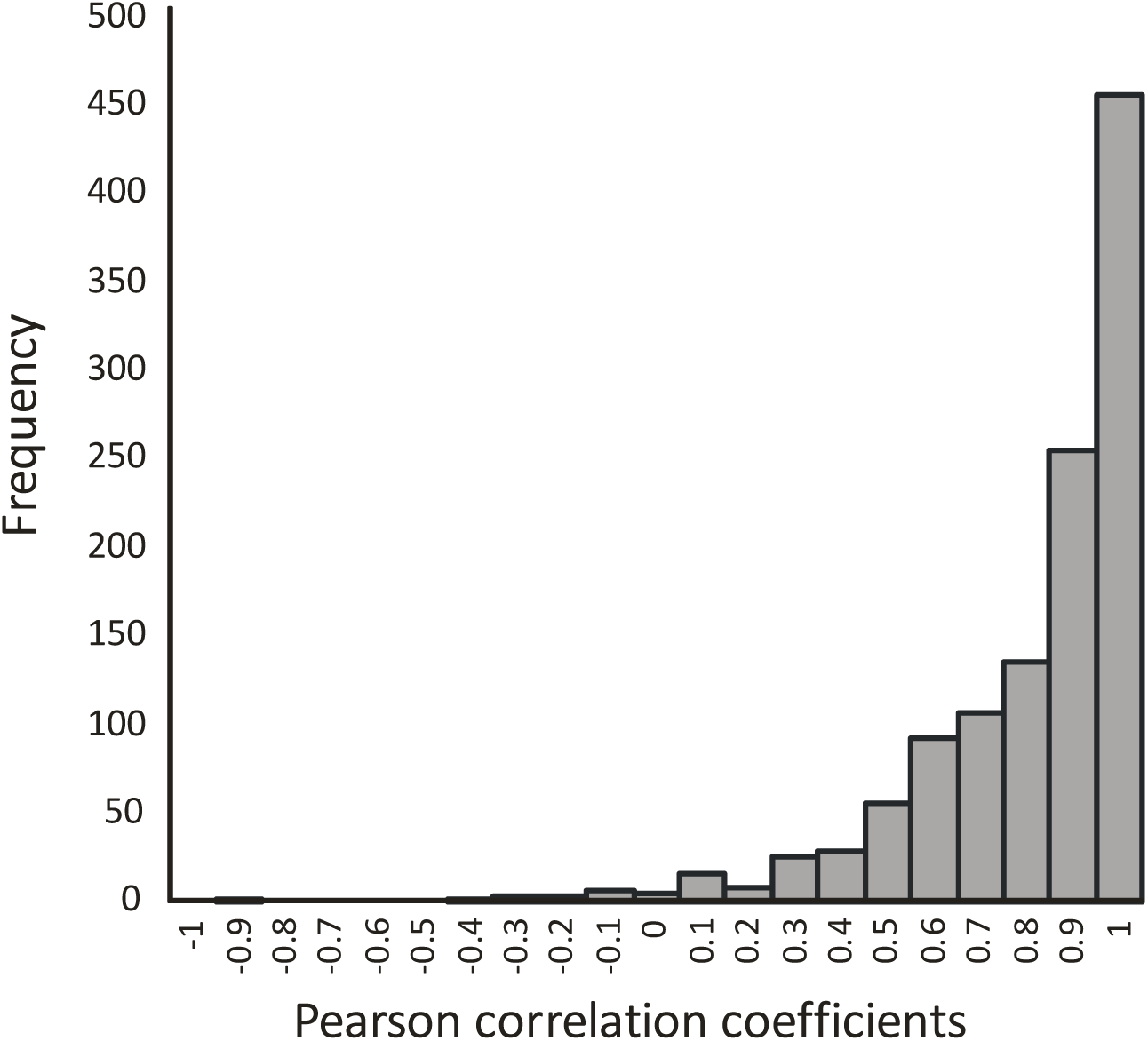
Distribution of correlation coefficients between the phenotypic effect of a QT locus and its observed allele frequency shift during adaptation.

Overall, almost half of the QTL loci changed their frequency significantly and could thus be detected (0.469, range 0–1). The most important significant factor here was N_b_ (70% variance explained), followed by the number of QTL (11%). Other significant factors had almost no (<1%) influence (Table 3). The detected loci explained almost three-quarters of the observed phenotypic change (0.738, range 0–1). The proportion of explained phenotypic change was by far best explained by N_b_ (78%). The remaining factor explained together less than 5% of variance (Table 4, Figure 3).

**Table 3.** Full GLM to explain variance in the percentage of detectable loci.

| Factors | d.f. | SS | MS | F | p | | $\eta^2$ |
| --- | --- | --- | --- | --- | --- | --- | --- |
| $N_b$ | 1 | 174 | 174 | 4960 | <2e-16 | *** | 0.701 |
| QTL | 2 | 28.2 | 14.1 | 399.9 | <2e-16 | *** | 0.113 |
| alpha | 1 | 0.03 | 0.03 | 0.814 | 0.367 |  |  |
| $x_{juv}$ | 1 | 1.65 | 1.65 | 46.83 | 1.24e-11 | *** | 0.007 |
| $N_b:QTL$ | 2 | 2.33 | 1.16 | 33.02 | 1.12e-14 | *** | 0.009 |
| $N_b:alpha$ | 1 | 0.01 | 0.01 | 0.373 | 0.541 | | |
| QTL:alpha | 2 | 0.42 | 0.21 | 5.974 | 0.003 | ** | 0.002 |
| $N_b:x_{juv}$ | 1 | 0.42 | 0.42 | 11.94 | 0.001 | *** | 0.002 |
| QTL:$x_{juv}$ | 2 | 0.02 | 0.01 | 0.303 | 0.739 | | |
| alpha:$x_{juv}$ | 1 | 0.00 | 0.00 | 0.121 | 0.728 | | |
| $N_b:QTL:alpha$ | 2 | 0.05 | 0.03 | 0.766 | 0.465 | | |
| $N_b:QTLi:x_{juv}$ | 2 | 0.02 | 0.01 | 0.320 | 0.726 | | |
| $N_b:alpha:x_{juv}$ | 1 | 0.00 | 0.00 | 0.025 | 0.874 | | |
| QTL:alpha:$x_{juv}$ | 2 | 0.02 | 0.01 | 0.349 | 0.705 | | |
| $N_b:QTL:alpha:x_{juv}$ | 2 | 0.02 | 0.01 | 0.320 | 0.726 | | |
| Residuals | 1176 | 41.45 | 0.04 |  |  |  |  |

**Table 4.** Simplified GLM to explain the percentage of phenotypic change explained by identified QTL.

| Factors | d.f | SS | MS | F | p | | $\eta^2$ |
| --- | --- | --- | --- | --- | --- | --- | --- |
| N<sub>b</sub> | 1 | 255 | 255 | 5562 | <2e-16 | *** | 0.788 |
| QTL | 2 | 11.6 | 5.78 | 126.3 | <2e-16 | *** | 0.036 |
| alpha | 1 | 0.42 | 0.42 | 9.099 | 0.003 | ** | 0.001 |
| X<sub>juv</sub> | 1 | 1.16 | 1.16 | 25.31 | 5.63e-07 | *** | 0.004 |
| N<sub>b</sub>:QTL | 2 | 0.46 | 0.23 | 4.989 | 0.007 | ** | 0.001 |
| N<sub>b</sub>:alpha | 1 | 0.08 | 0.08 | 1.853 | 0.173 |  |  |
| QTL:alpha | 2 | 0.11 | 0.06 | 1.249 | 0.287 |  |  |
| N<sub>b</sub>:QTL:alpha | 2 | 0.30 | 0.15 | 3.241 | 0.039 | * | 0.001 |
| Residuals | 1187 | 54.4 | 0.05 |  |  |  |  |

**Figure 3.**
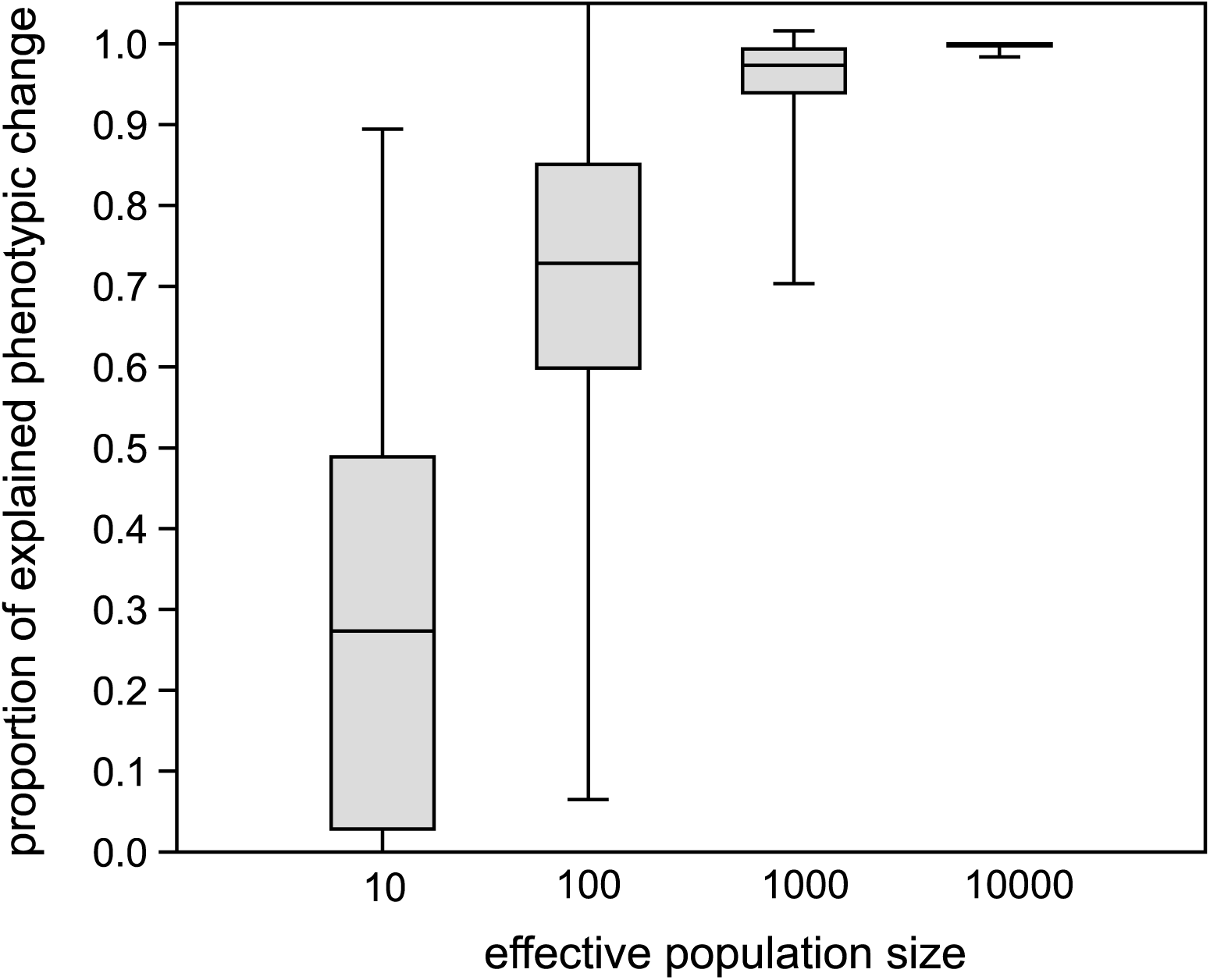
Boxplot of the effect of number breeding adults on the proportion of explained phenotypic change by detectable loci.

## Discussion

In the simulations, the speed of adaptation depended largely on the number of offspring per breeding pair, as initially expected. Notably, this not an effect of the total number of offspring per generation. When comparing simulations with the same total number of offspring (e.g. N_b_ 100, 200 x_juv_ and N_e_ 10000, 2 x_juv_), always the scenario with more offspring per breeding pair evolves faster. Overall, adaptation proceeded with remarkable speed in terms of generations (Figure 1). The relative independence from the number of breeding pairs suggested that even small populations can evolve efficiently and comparatively rapidly at least in quantitative traits as long as there is enough standing genetic variation. The genetic architecture in terms of number of loci involved and differential contribution distributions of individual loci (alpha) was found rather unimportant for the adaptation speed. Large offspring numbers may thus at least partially explain the increasingly observed instances of rapid adaptation.

Whether the presented results are indeed relevant in nature depends on the prevalence of the chosen deterministic soft selection regime. It appears that density and/or frequency dependent selection regimes, where the individuals closest to some new or spatially and/or temporally varying trait optimum survive or get to reproduction with higher probability are the most frequent in nature [11]. However, less deterministic scenarios than the one used here with more random components are likely more realistic, but they also introduce more noise, thus blurring the possibility to infer the influence of the investigated factors. It is the particular strength of simulations to simplify assumptions to make predictions, explore processes or develop new theories [14]. Moreover, the chosen deterministic selection is directly relevant at least for evolutionary experiments [15] and artificial selection [16].

The suggested effect of offspring numbers on adaptation speed may also contribute to explain variance of offspring numbers among evolutionary lineages. In populations at carrying capacity and thus constant population size, any given breeding pair will contribute on average two offspring to the next breeding generation. Gross overproduction appears counterintuitive in crowding situations as more investment in less offspring should be expected [17] and has been shown empirically [18].

However, if the environment of a population is highly variable and unpredictable, a large number of offspring may be a genetic strategy in the first place. A large number of offspring fills the possible phenotype space as determined by the genotypes of the parents for the quantitative trait in question as broadly as possible (Suppl. Fig. 2). A large number of offspring thus increases the chances to produce at least a few close to the (unpredictable) fitness optimum in the next generation. Producing much more offspring than can possibly survive may be therefore not a waste of resources but rather a sort of genetic bet hedging strategy. I found here that maximum adaptive speed was already reached with a few dozen offspring. This seems to contradict the often much higher observed offspring number per pair in nature [19]. However, if, as may be expected in nature, more than a single quantitative trait needs optimisation under changing ecological conditions, respectively higher offspring numbers may be necessary to achieve fast adaptation in all of these traits simultaneously.

The results on the detectability of the loci underlying the observed phenotypic change with population genomic scans from before-after data are encouraging (Table 3). As might have been expected, detectability depended largely on the number of breeding adults, i.e. a proxy for the effective population size. The larger the breeding population, the more negligible allele frequency changes due to genetic drift became at neutral loci. Before such a background, the (larger) allele frequency shifts due to selection stood out and could be statistically identified (Suppl. Fig. 1). Whether the observed moderate drift at neutral loci is only due to the effective population size as determined by the number of breeding adults or whether increased variance in offspring due to selection additionally played a role, remains to be studied. However, other selection scenarios e.g. on differential fecundity (winner-takes-it-all scenario) rather than survival, would increase the variance in offspring much more and thus reduce the effective population size and increase drift [20]. Depending on the loss of genetic variation due to this drift, adaptation is perhaps slowed down or even prevented. Such scenario could also be tested with the IBM model presented here. Interestingly, there was generally a strong correlation between the magnitude of effect a locus had on the phenotypic trait and its response to selection (Fig. 2), as expected [6]. However, in a considerable fraction of simulations, this correlation was weak or even absent (Fig. 2).

Already with moderate breeding population sizes (100 individuals), the large majority of the loci responsible for the observed phenotypic change could be identified and an even larger fraction of the observed phenotypic change accounted for (Table 3). This makes experimental approaches to test the hypothesis outlined here feasible and promising.

## Acknowledgements

I am grateful to Barbara Feldmeyer for help with the GLM.

